# Decision Support Systems based on Scientific Evidence: Bibliometric Networks of Invasive *Lantana camara*

**DOI:** 10.1101/2020.08.10.240879

**Authors:** Preet Mishra, Abhishek Prasad, Suresh Babu, Gitanjali Yadav

## Abstract

Extraction and analysis of useful knowledge from the vast amount of relevant published literature can add valuable insights to any research theme or area of interest. We introduce a simplified bibliometric data analysis protocol for gaining substantial insights into research thematics, which can also serve as a handy practical skill for researchers, while working from home. In this paper, we provide ways of developing a holistic research strategy using bibliometric-data driven approaches that integrate network analysis and information management, without the need of full paper access. This protocol is a comprehensive multi-modular pathway for analysis of metadata obtained from major scientific publishing houses by use of a Decision Support System (DSS). A simple case study on the invasive species *Lantana camara* has been presented as a proof-of-concept to show how one can implement this DSS based protocol. Some perspectives are also provided on how the outcomes can be used directly or scaled up for long term research interventions. We hope that this work will simplify exploratory literature review, and enable rational design of research objectives for scholars, as well as development of comprehensive grant proposals that address gaps in research.

## INTRODUCTION

### The problem of optimizing knowledge extraction from published literature

Informed policy making necessitates scientific evidence, regardless of system complexity, be it socio-political state priorities during the ongoing novel coronavirus pandemic, funding agency mandates, research strategy development of graduate students, or any other real world problem. While identifying possible solutions, policy makers require problem-oriented scientific evidence, which in turn, can be understood as a dynamical system possessing a multi-modular nature. Accordingly, policy issues and solutions are both embedded in complex, globally interconnected environments that require thorough analysis of knowledge extracted from available published or bibliometric data. Any evidence derived from incomplete data can be inaccurate, and this brings in two critical aspects associated with usage of vast information:

Firstly, published information can be categorized under the paradigm of big data, possessing all the typical characteristics of high variety and volume. Usage of such a large body of information is a complex process. Thus, the analysis of information, if not implemented without due recognition of constraints involved, such as copyright, open access, metadata etc. can cause bottlenecks in data adequacy.

Secondly, data insufficiency may arise from the choice of search parameters. Computational tools like citation managers and AI-based search engines harness bibliometric big-data using author-designated and user-specified “keywords”. This step, in turn, controls all the later stages of the analysis; organizing, filtering, classifying, predicting, planning and so on. If a priori, the keyword is chosen in an ad-hoc or static manner, it may decrease efficiency of the search process which in turn will strongly impact outcome.

Since both these aspects of the problem are inherent and user-dependent, we suggest a two-pronged strategy of building redundancies and optimizations using decision support systems, as described below.

### Visualizing bibliography metadata as Networks

Network approach to decipher large data sets has long been known as one of the best analytical strategies, and this applies equally well for bibliographic information (Shiffrin and Börner, 2004), with the added benefits of being able to explore, model and restructure literature metadata to draw insights from both static and dynamic representations of individuals, organizations or themes of research (Newman, 2004). High dimensional literature metadata, when visualized efficiently through networks can reveal communities sharing common node or edge attributes in both coarse-grained and fine-grained routines (Babu *et al*., 2016)

Availability of metadata can often overcome constraints of limited access to full-text and enable one to focus on a lower ease-of-access threshold, i.e. critical information within the title, abstract and citation. In the paradigm of embedding theory, words embedded within sentences of publications indicate the themes of research and thus the visual analysis of these embeddings may provide proof of concept (Spangler *et al*., 2014). Such textual embedding of data has high dimensions (Griffiths and Steyvers, 2004) and keeping track of the dimensions can be done in an efficient manner through network visualizations of co-occurrences (Tshitoyan *et al*., 2019). An added benefit is, it can provide us a visual static picture of the flow of ideas and the themes in current research scenarios (Srinivasan, 2004). Metadata can thus consist of keywords, title, and abstract, authors’ information, publisher and journal information which can all be projected as node attributes, while edges may represent relationships between nodes, such as co-authorship, professional affiliations, or collaborative interactions (Landauer *et al*., 2004). Co-authorship bibliography networks are undirected, and two of the most informative topological parameters are (a) betweenness centrality, an indicator of hubs in the flow of information, and (b) degree distribution, a measure of the collaborations (Newman *et al*., 2004).

### Decision support system (DSS)

A vast number of online and downloadable tools are available to perform bibliometric analyses, but as mentioned above, all such software strictly adhere to the concept of Garbage In Garbage out (GIGO), being implementations based on the processing of metadata obtained through user-derived “keywords” (Krallinger *et al*., 2017). If we envisage it a single process to obtain metadata from a huge database of published articles, then it may be possible to optimize critical search parameters, through a Decision Support System (DSS) (Sprague, 1980).

Here we present case studies from invasive plant species bibliographic metadata and share how emerging co-authorship networks can improve and inform decisions, and how diverse network visualizations can be integrated as modules in a DSS. The solution provided from these DSSs must be regarded as partial, being iterative and adaptive, subject to existing constraints in time, rather than being steadfast or all encompassing. Our main objective is to increase the efficiency of decision making at the initial levels of strategy design, which may contribute to accuracy improvement at later levels in the hierarchy.

## METHODLOGY

### Design of the DSS

Figure 1 depicts a schematic of the suggested decision support system (DSS) for integration of modules consisting of various bibliography tools, to analyze the metadata of published information.

**Figure 1.**
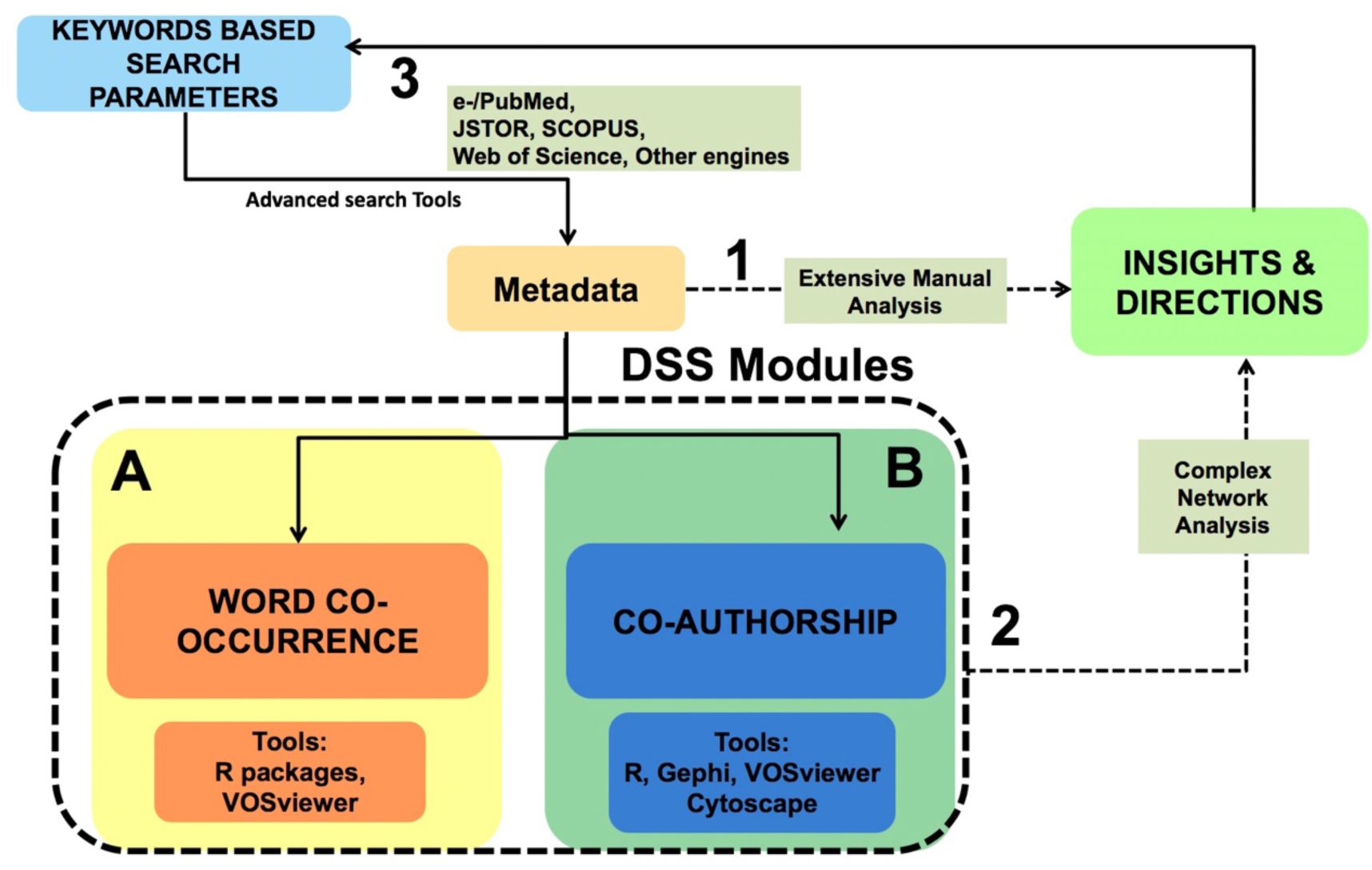
DSS Schematic. Routes 1 and 2 mark options to assess bibliometric metadata, with the latter involving modular analysis of bibliometric networks (A,B) as a Decision Support System.

A-priori the user-selected keyword enters the process, and is at the core of advanced search tools for obtaining metadata from various sources like. This metadata can be analyzed in two distinct ways (dashed line arrows marked as 1 and 2 in Fig 1); route 1 includes labor-intensive reading and analysis of the problem-oriented search results and then updating, on a manual inference basis, the search keyword in the next iteration, a complex and multi-dimensional landscape that may involve new queries arising from parsing of metadata and analytical constraints related to accessibility, time availability, feasibility, collaborations, and so on. This juncture can be identified as the most crucial point in the analysis, where user has to make decisions regarding the next iteration. Several queries need to be addressed before moving ahead in an exploratory investigation.

For instance, how much information should be sufficient to answer the questions addressed? Do the current citations cover adequate thematic premise? Does the metadata reflect evolution of subject-specific knowledge through space and time? According to our proposed scheme, route 2 can be used to answer these and other relevant questions through a DSS with two modules A and B, both incorporating diverse aspects of network theory (See Figure 1).

### Bibliography Data Formats

Article metadata can provide information at the theme levels and thus it can be related to decisions about choice of keywords and databases to search when doing the literature surveys and concept reviews broadly. Module A in Figure 1 involves text co-occurrence networks based on the metadata obtained from initial searches. These visualizations can be obtained from the abstract, title, or keywords provided with the published paper. Some of the file formats that are typically used are .txt, .csv, .xml etc. These can then be plugged into the respective tools to obtain relevant visualizations. Some of the tools are mentioned in the Figure 1. Through various algorithms, these tools parse the embeddings of the words in the text and yield mapping of the embeddings, providing users with a clearer picture of the linked concepts, which have been published and latent information which can help users select appropriate keywords for the next iteration.

The metadata obtained from the search can also yield information about the research scenarios at the author level with related information as attributes of the author. The module marked B in Figure 1 deals with user decisions regarding the choice of the search parameters taking into account author based information from metadata. This module involves tools to create co-authorship networks that can be analyzed regarding not-only decisions about search parameters but also inform the exploratory investigation into research themes by answering questions as to the origins and extent of the problem itself, i.e. who, where, what and when etc. (Lent *et al*., 1997). Both DSS modules are highly user-centric and involve open source programming software, with greater levels of accessibility, enabling personalized module design at user level. Module B depends on the interpretation of the user in a more convoluted way than that involved in module A, and due care must be taken while implementing module B, as well as while deriving interpretations from the DSS outputs at each iteration.

## RESULTS

### A case study on *Lantana camara*

We present a case study for assisting a graduate student aiming to perform a systematic review of biological invasions, focusing on India’s most aggressive alien plant invasive species *Lantana camara*. Our objective is to roughly get an overview of the status of research in this area and identify major gaps, specially from an Indian context, so that a suitable graduate research strategy can be designed for the next three years, addressing gaps in knowledge.

For this case study, we take route 2 in Figure 1 and show how combinations of suitable keywords can be developed by multiple iterations of the DSS module, the methodology is summarized as a flowchart in Figure 2.

**Figure 2.**
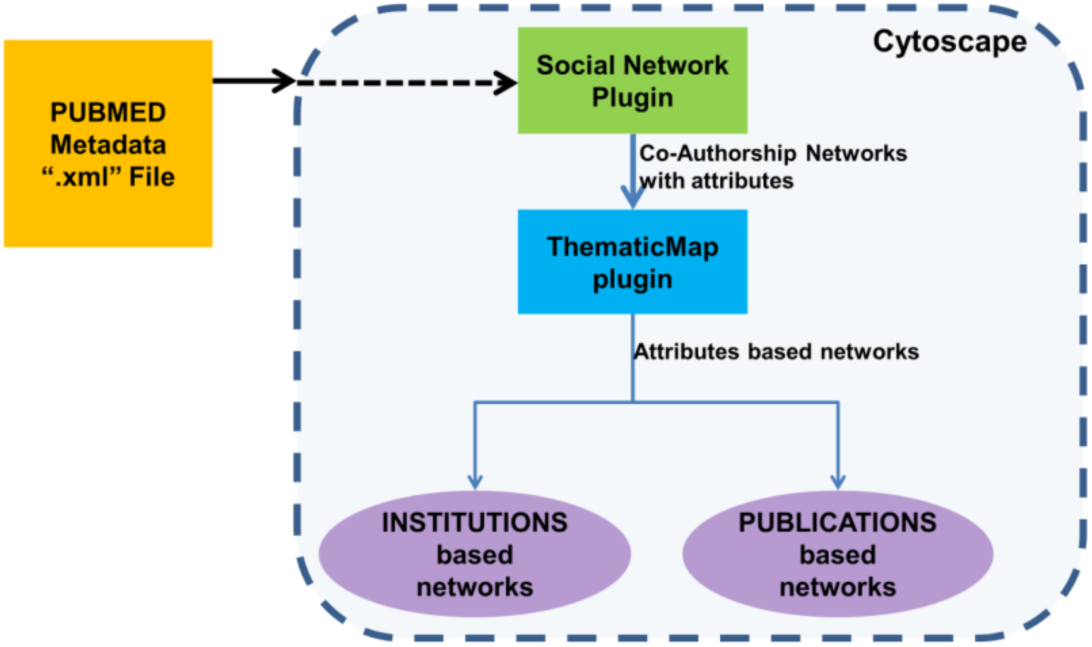
The flow chart of the methodology adopted for the case study.

For the initial process of obtaining metadata, we focus on the prime objectives of the species in Indian context. We use PUBMED (https://www.ncbi.nlm.nih.gov/pubmed), the open-source literature database with over 30 million citations for biomedical literature beginning from 1948, life science journals, and online books (https://www.ncbi.nlm.nih.gov/books/NBK3827). We used ‘advanced search’ option and keywords (“lantana”[All Fields]) AND (“camara”[All Fields]) AND (“India”[All Fields]), and this returns over a 100 articles by more than 300 authors. We saved metadata as an XML file, a format compatible with the Cytoscape plugin used in subsequent stages (Shannon *et. al*., 2003). It may be noted that formats and compatibility between search engines and downstream tools represents a constraint in the visualization process and metadata formats with more flexibility can enable more information to be extracted. For instance, although JSTOR (www.jstor.org) has the largest open collection of published articles (1928 onwards), it does not enable metadata collection, rather each article is downloaded as an individual XML file. We are currently working on developing a tool for collating these files and extract metadata using R without losing links between the metadata. It may also be noted that we have chosen a relatively sparse dataset as our example, and we advise due care in selection of keywords and to correlate these with number of articles returned. For example, adding the keyword ‘invasive’ severely reduces the number of articles, whereas removing the connection to India increases the number to beyond 600.

The XML file feeds into the ‘Social Network’ plug-in of the most widely used open source visualization tool Cytoscape (Kofia *et al*,. 2015), yielding the co-authorship bibliography network shown in Figure 3A. Various attributes of the nodes (authors) can be superimposed onto this network as additional informative layers of color or shape, such as for example, the institutions they are affiliated to, as well as theme of research, to obtain visualizations in panels B and C respectively (Figure 3).

**Figure 3.**
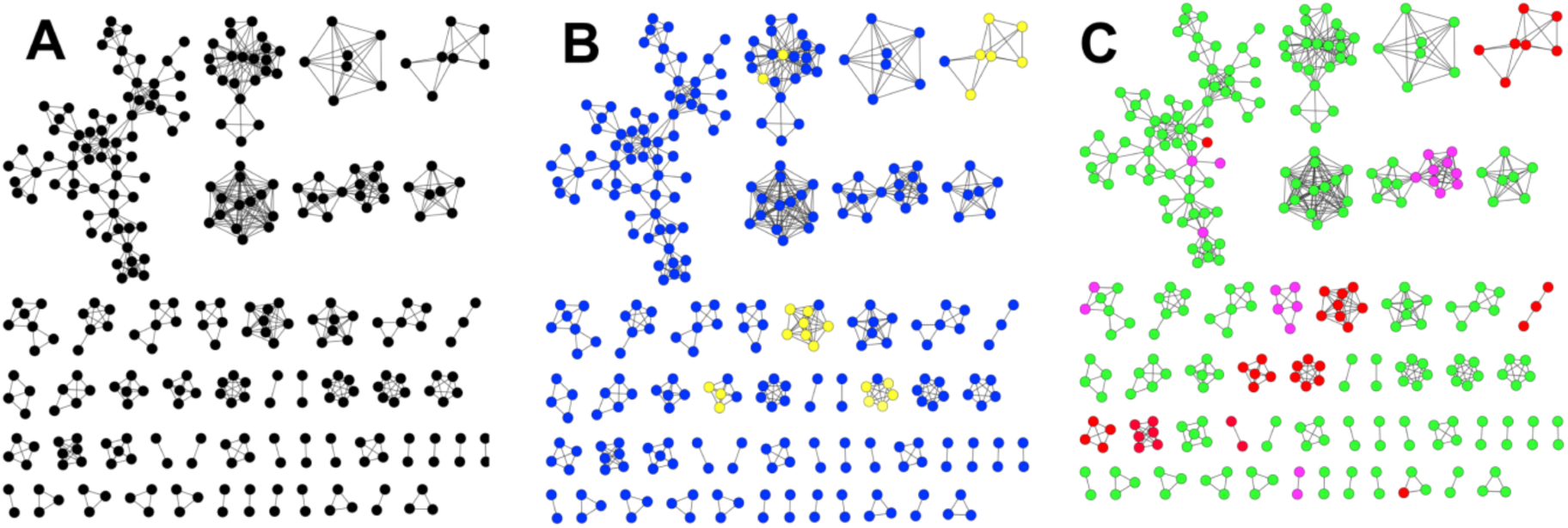
Co-Authorship Networks overlaid with metadata (A) Nodes are authors and edges represent shared publications. (B) Same as A but color-coded by author affiliations; Blue nodes depict authors from Indian Institutions. (C) Same as A but color coded by themes/keywords; 42 Reds (Invasion Ecology), 249 Greens (Applied Phytochemistry) and 15 Pink nodes (depicting agricultural applications) in relation to *Lantana camara*.

The foremost observed pattern is that Lantana research in India is strongly inclined towards applied phytochemistry, with more than 80% of authors publishing under this theme (represented by green colored nodes in Fig 5C). The other notable pattern is that despite very few researchers (only 13%), being involved in addressing invasion ecology of *L*.*camara* in India this is the theme where potential global interest, and possibly funding opportunities, may be found. This is evident from a comparison of congruent layouts in Figures 5B and 5C, where largest proportion of foreign authors (yellow nodes in 5B) are sharing publications with Indian authors under the theme of invasion ecology (represented by red nodes in 5C).

In order to check if this interpretation is an artefact of network representation, the layouts can be redrawn differently; representing nodes as institutions or publications, rather than individual authors. This has been done in Figure 4 using the same metadata and color codes as above, with Cytoscape ‘ThematicMap’ plugin (Shannon *et. al*., 2003). All attribute networks in Figure 4 represent edges as authors and the width of these edges depicts the number of authors sharing an affiliation (4A, 4B) or a publication (4C). Colors in Figure 4A reveal collaborations between Indian (blue) and foreign (yellow) institutions, while color codes in Figure 4C reveal theme of the paper. Clearly, the pattern observed in the earlier networks is echoed here as well, with 80% of the network space representing applied phytochemistry, with the added benefit of being able to map organisations to research themes of interest, such as invasion aspect *Lantana camara*.

**Figure 4.**
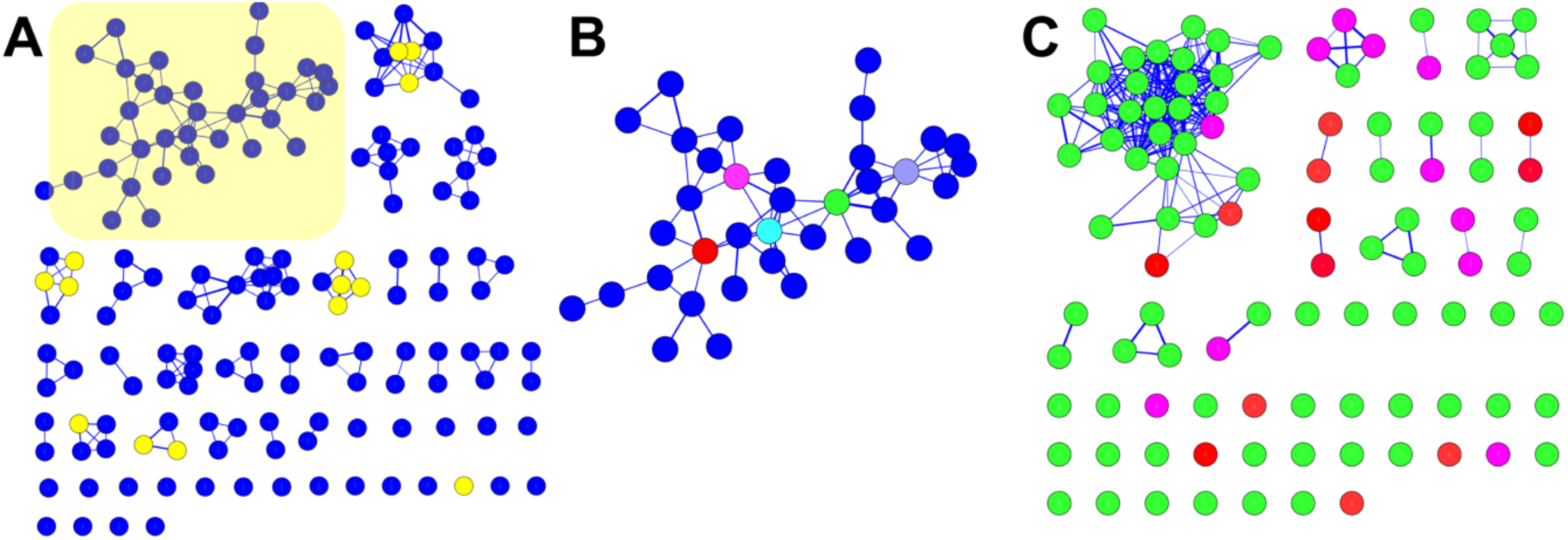
Attribute networks for *L*.*camara* literature (A) Nodes are Indian (blue) or foreign (yellow) institutions while edges represent the number of authors sharing the affiliation. (B) Largest subnetwork highlighted in 4A, Top 5 institutes with highest topological centralities are depicted in new colors (see text). (C) Nodes are papers and the edges represent common authors. Color codes same as in Figure 3.

Additional features of the bibliography networks can be observed in attribute networks, as compared to co-authorship networks; Figure 4B shows the largest subnetwork of 4A, revealing the most highly associated institutions among existing collaborations, and this subnetwork should be important for decisions regarding setting up new collaborations, or identification of hubs driving the existing body of work, towards bringing in fresh outlook or scope. Nodes in the 4B subnetwork are colored by degree, revealing institutions that have had the maximum collaborative interaction in the area of interest. Interestingly, this subnetwork consists of organisations that are not only related by geographical proximity (like IVRI and CSIR-IHBT both in Palampur, and Doon University, Dehradun), but also connect a wide diversity of expertise (Dept of Biochemistry at Panjab University and Carbohydrate Biotechnology at IIT Guwahati), both being valuable for a new researcher.

Additional features discernable from both sets of networks are related to linkages of research and development units, towards reduction of multidimensional problems related to research in invasion biology of *Lantana camara*. For instance, agricultural applications are not very strongly represented in Lantana bibliography networks (only 4%), but this is not a niche or gap in research, instead, a reflection of the nature of *Lantana camara;* it is not an agricultural weed but more widely impacting the conserved/protected areas. As can be seen in Figure 3C and 4C, the few studies that involve agricultural applications, are also in the area of applied photochemistry, suggesting an applicability for Lantana phytochemicals in toxicity assays or weedicide development. The major research gaps from an Indian point of view in this field are that ecology and phytochemistry have not been studied together for this invasive, despite being known to be connected concepts. Another insight from these networks is the large number of disconnected subnetworks, suggesting a relative disconnect between researchers, groups and institutions working on *Lantana camara*. This awareness immediately raises the need for a pan-India national network on invasion biology, bridging diverse fields of expertise and knowledge, to address this most aggressive species that has invaded almost all parts of the country.

Among the keywords that may enable more detailed literature review and decision making, would be to expand the search to larger engines like Web of Science and JSTOR, as well as to get a global perspective by dropping the reference to India alone. Web of Science has over 1000 articles while JSTOR has over 2000 articles in this category. We have combined both searches and are presently developing R scripts to link the metadata from all sources for a more comprehensive and global analysis. Another option is to perform the next iteration of this DSS by adding more specific theme-specific keywords, phytochemistry being of foremost importance. With the largest number of paper available, new questions could be addressed in phytochemical evolution. Another immediate consequence of this small study would be to expand the search to other invasive alien species, an effort that has already been undertaken in our group. The metadata can also be analyzed using co-occurrence visualization software like VOSviewer (van Eck and Waltman, 2010), that forms the other module of the proposed DSS (module B; Figure 1).

## OUTLOOK AND PROSPECTS

In this work, we have shown that even with a simple DSS on a single iteration, the outcome not only enabled identification and optimization of multiple relevant keywords, but also put forth a complete picture of the various aspects of the research theme, that emerged outright at the initial stages itself. We also provide ways to scale up and the search operation with additional databases, as well as scaling down with selection of a range of keywords.

Two courses of action emerge from the DSS and case study presented here. First, a GUI tool could be designed with integrated modules of the above visualization-based analysis as separate programs, which when executed by the user, would act as a source of both quantitative and pictorial depictions of results (Skupin, 2004). The process of refining the input to such a tool could then be performed iteratively to obtain precise insights at each step of the procedure. Higher levels of categorization tools exist beyond a high threshold of ease-of-access thus, open-access tools are needed to tackle the issues of a good literature review for usage by a wide spectrum of researchers.

Second, individual-based support system can be designed subject to the constraint of know-how and ease of access to the above-mentioned network visualizations tools. For example, searching of metadata from web based databases can be scripted through text and data mining tools like ContentMine, and custom-built in open source platforms like R or Python. Combined with this, the knowledge of advanced network analysis tools like Gephi, Pajek etc. can be used then to analyse the metadata from various perspectives. This will help gather information that would assist the individual to make a holistic research strategy.

In summary, we hope that our work will pave the way for new scientific-evidence based insights, policy decisions and future directions through streamlined network analytics. In exceptional circumstances such as the present Covid-19 lockdowns, a lot of review work and metanalysis is being carried out globally. With a few strategic interventions, many of these *in silico* efforts could provide insights into gaps and opportunities in research thematic across a range of disciplines.

## Author Contributions

SB and GY designed the study and proof of concept. PM and AP collected the data. PM built the DSS and performed the analysis. All authors contributed to preparation of the manuscript.

## Acknowledgements

AP acknowledges CSIR Junior Research Fellowship. GY acknowledges SERB grant ID EMR/2016/006486, GCRF-BBSRC grant ID BBSRC BB/P027970/1TIGR2ESS (Transforming India’s Green Revolution by Research and Empowerment for Sustainable food Supplies), and NIPGR support. PM acknowledges JNU and UGC for support. SB acknowledges support from AUD and CUES during PM’s internship for this work.

## REFERENCES

1. Babu S and Yadav G (2016) Co-Authorship Networks among DRDO Life Science Scientists Defence Life Science Journal 1 188–191

2. Griffiths T L and Steyvers M (2004) Finding scientific topics Proc. National Academy of Sciences 101 (suppl 1) 5228–5235

3. Kofia V, Isserlin R, Buchan A M and Bader G D (2015). Social Network: a Cytoscape app for visualizing co-authorship networks. F1000Research 4 481.

4. Krallinger M, Rabal O, Lourenço A, Oyarzabal J & Valencia A (2017) Information retrieval and text mining technologies for chemistry Chem. Rev. 117 7673–7761.

5. Landauer T K, Laham D, Derr M (2004) From paragraph to graph: Latent semantic analysis for information visualization Proc. National Academy of Sciences 101 (suppl 1) 5214-5219

6. Lent B, Agrawal R, and Srikant R (1997) Discovering trends in text databases In Proceedings of KDD, International Conference on Knowledge Discovery, NewPort Beach CA 227– 230.

7. Newman M E J (2004) Coauthorship networks and patterns of scientific collaboration Proc. National Academy of Sciences 101(suppl 1) 5200–5205.

8. Newman M E J and Girvan M (2004) Finding and evaluating community structure in networks Phys. Rev. E 69 026113

9. Shannon P, Markiel A, Ozier O, Baliga N S, Wang J T, Ramage D, Amin N, Schwikowski B and Ideker T (2003) Cytoscape: a software environment for integrated models of biomolecular interaction networks Genome Research 13 2498–504

10. Shiffrin R M and Börner K (2004) Mapping knowledge domains Proc. National Academy of Sciences 101(suppl 1) 5183–5185

11. Skupin A(2004) The world of geography: Visualizing a knowledge domain with cartographic means Proc. National Academy of Sciences 101(suppl 1) 5274–5278

12. Sprague RH (1980) A Framework for the Development of Decision Support Systems MIS Quarterly 4 1–26

13. Spangler S, Wilkins AD, Bachman BJ, Nagarajan M, Dayaram T, Haas P, Regenbogen S, Pickering CR, Comer A, Myers JN, Stanoi I, Kato L, Lelescu A, Labrie JJ, Parikh N, Lisewski AM, Donehower L, Chen Y and Lichtarge O (2014) Automated hypothesis generation based on mining scientific literature In Proc. 20th ACM SIGKDD Intl Conf. Knowledge Discovery and Data Mining 1877–1886

14. Srinivasan P (2004) Text mining: Generating hypotheses from MEDLINE J. Am. Soc. Inf. Sci. 55 396–413

15. Tshitoyan V, Dagdelen J, Weston L, Dunn A, Rong Z, Kononova O, Persson KA, Ceder G and Jain A (2019) Unsupervised word embeddings capture latent knowledge from materials science literature. Nature 571 95–98

16. van Eck, NJ, Waltman, L (2010) VOSviewer, a computer program for bibliometric mapping Scientometrics 84 523–538

